# Impact of Obesity on the CCR6-CCL20 Axis in Epidermal γδ T Cells and IL-17A Production in Murine Wound Healing and Psoriasis

**DOI:** 10.1101/2024.04.09.588780

**Authors:** William Lawler, Tanya Castellanos, Emma Engel, Cristian R. Alvizo, Antolette Kasler, Savannah Bshara-Corson, Julie M. Jameson

## Abstract

Obesity is associated with comorbidities including type 2 diabetes, chronic nonhealing wounds and psoriasis. Normally skin homeostasis and repair is regulated through the production of cytokines and growth factors derived from skin-resident cells including epidermal γδ T cells. However epidermal γδ T cells exhibit reduced proliferation and defective growth factor and cytokine production during obesity and type 2 diabetes. One of the genes modulated in epidermal γδ T cells during obesity and type 2 diabetes is CCR6, which is the receptor for CCL20. CCL20 is elevated in the skin during obesity and type 2 diabetes. Here we identify a subset of murine epidermal γδ T cells that expresses CCR6 in response to activation *in vitro* and post-wounding or psoriasis induction with imiquimod *in vivo*. We show that CCL20 stimulates epidermal γδ T cells to produce IL-17 suggesting CCR6 regulates the IL-17 axis as in dermal γδ T cells. Further, epidermal γδ T cells upregulate CCR6 and produce IL-17 during murine models of wound repair and psoriasis. Obesity increases CCR6 and IL-17 expression by epidermal γδ T cells during wound repair but has less of an effect during psoriasis. These findings have novel implications for the regulation of a specific population of IL-17-producing epidermal γδ T cells during skin damage and inflammation.

## Introduction

Obesity is correlated with increases in skin comorbidities including chronic nonhealing wounds and psoriasis (1–4). Chronic nonhealing wounds affect 2.5% of the US population while psoriasis affects 8 million Americans (1, 5). Normally the skin provides a protective barrier from mechanical, chemical, and pathogenic external threats. The TNF-α-IL-17 axis provides antimicrobial roles and induces keratinocyte proliferation and neutrophil responses in wound repair (6, 7). However, dysregulation of the TNF-α-IL-17 axis caused by obesity alters the cellular composition and function in the skin resulting in premature keratinocyte differentiation, altered skin-resident T cell number and function, and increased barrier permeability (7, 8). All of these factors negatively impact chronic nonhealing wounds and psoriasis (1).

Alterations in TNF-α and IL-17 production during chronic nonhealing wounds and psoriasis have been attributed to dermal γδ T cells, dermal Th17 αβ T cells, innate lymphoid cells (ILC), and mucosal-associated invariant T (MAIT) cells (9–12). In contrast, resident epidermal γδ T cells, also known as dendritic epidermal T cells (DETC), have been largely considered bystanders and not active IL-17 producers (13). Epidermal γδ T cells bear the Vγ5Vδ1 TCR and are rapidly activated by stressed or damaged keratinocytes to release cytokines, chemokines, and growth factors including TNF-α and in some cases IL-17A (14–16). In lean mice, epidermal γδ T cells serve as critical players in skin homeostasis and wound healing (17–20). However, obesity causes a reduction in TNF-α production by epidermal γδ T cells at the wound site indicating a shift in function (15). Since epidermal γδ T cells normally act early in wound repair, they may also negatively impact chronic wounds and inflammatory skin disease (13).

One receptor that has been associated with IL-17 producing dermal γδ T cells (Tγδ17) is CCR6. The majority of murine dermal Vγ4 and Vγ6 T cells express CCR6, which facilitates recruitment to the epidermis in response to psoriasis-like inflammation (13, 21, 22). CCR6 is also required for efficient wound repair as CCR6^−/−^ mice exhibit delays in wound closure (23). Obese and diabetic patients exhibit elevated CCL20 in the skin, which likely impacts CCR6^+^ T cell recruitment and function(24). Although CCR6 regulates other IL-17-producing T cell populations, epidermal γδ T cells have not been included as they already reside in the epidermis and do not express CCR6 constitutively (13, 25). Thus, epidermal γδ T cells may exhibit unique regulation and function of CCR6 especially in obesity.

In this study we identify how epidermal γδ T cells participate in obesity-related skin complications such as wound healing and psoriasis-like inflammation. We find that CCR6 is expressed by a distinct population of activated epidermal γδ T cells and CCL20 induces this subset to produce IL-17. Using previously published single cell RNA sequencing (scRNAseq) data, we show that CCR6^+^ epidermal γδ T cells express IL-17-associated genes during psoriasis-like inflammation. We validate these findings in murine models of wound repair and IMQ-induced psoriasis-like inflammation where CCR6 and/or IL17A are expressed by epidermal γδ T cells *in vivo*. Obesity increases the number of epidermal γδ T cells expressing CCR6 and IL-17A during wound healing, which underscores the significant impact of obesity on skewing toward an IL-17 proinflammatory response.

## Materials and Methods

### Mice

C57BL/6N mice were purchased from Taconic Biosciences (Rensselaer, NY). 16 to 20-week-old male mice were studied for all cell culture and flow cytometry experiments. Male C57BL/6-Il17atm1Bcgen/J mice (JAX stock 018472) were purchased from The Jackson Laboratory (Bar Harbor, Maine). All C57BL/6-Il17atm1Bcgen/J mice used for experiments were purchased between the ages of 4-6 weeks of age and used between 18-24 weeks of age. Mice received access to food and water ad libitum and were housed in the animal facility at California State University San Marcos. For obesity studies, 6 week old male C57BL/6-Il17atm1Bcgen/J mice were fed either a high fat diet (HFD) consisting of 60 kcal% fat diet (D12492, Research Diets) or normal chow diet (NCD) (D12450J, Research Diets) for 12-16 weeks. All experimental procedures involving animals were reviewed and approved by the Institutional Animal Care and Use Committee of California State University San Marcos (21–003).

### Epidermal T Cell Culture

Epidermal cells were harvested from the back skin of 16 to 20-week-old wild-type mice as previously described (26). Briefly, the back skin was removed, cut into 1cm^2^ squares, and incubated on 0.3% trypsin-GNK (0.09% glucose, 0.84% sodium chloride and 0.04% potassium chloride) at 37°C with 5.0% CO_2_ for 3.5 hours. The epidermis was then peeled from the dermis and shaken in 0.3% trypsin GNK with 0.1% DNase at 37°C for 10 minutes. The cell solution was placed in DMEM with 10% heat inactivated FBS, 2.5% HEPES buffer, 1% 100x non-essential amino acids (NEAA), 1% 100mM sodium pyruvate, 1% penicillin streptomycin glutamine (PSG), vitamins and 0.1% 2-mercaptoethanol (2-ME). The cells were filtered through Sera-Separa filter columns, pelleted, and purified with Lympholyte M prior to culture. The epidermal cells were plated in a 96-well plate containing RPMI-1640 media with 10% FBS, 2.5% HEPES buffer, 1% NEAA, 1% sodium pyruvate, 1% PSG, 0.1% 2-ME, and 20 U/ml Interleukin 2 (IL-2). 2μg/ml of Concanavalin A (ConA) and 1μg/ml of indomethacin were added at the initiation of cell culture. Twice a week fresh media without ConA and indomethacin was added. Cells were restimulated every 3 weeks with 1μg/ml of ConA and harvested when epidermal γδ T cells made up over 95% the cells in culture (typically 10 weeks).

### Epidermal γδ T Cell Activation

For *in vitro* studies, 24-well plates were precoated with 1μg/ml or 10μg/ml of anti-CD3ε for 24 hours at 37°C with 5.0% CO_2_. The supernatant was harvested and stored at −80°C for Luminex analysis. For intracellular cytokine staining, the cells were stimulated for 6 hours and 5ug/ml Brefeldin A (BFA) was added at hour 2. BD fixation and permeabilization kit (BD Cytofix/Cytoperm) was used per manufacturer’s instructions. The following antibodies were used: anti-CD3 (145-2C11), γδ TCR (GL3), CD25 (3C7) and CCR6 (29-2L17) (Biolegend, San Diego, CA). Flow cytometry was performed on a Accuri C6 (BD Biosciences, San Diego, CA) and data was analyzed with FlowJo software (BD Biosciences).

### IMQ Induction of Psoriasis-like Inflammation

5mg of 5% IMQ cream (IMQ Cream 5%, Fougera Pharmaceuticals Inc.) was applied to the right ear and control cream (Vanicream, Pharmaceutical Specialties, Inc.) applied to the left ear of C57BL/6-Il17atm1Bcgen/J mice daily for up to 3 days (9). Mice were individually housed and monitored daily. At the end of the experiment, ears were harvested for staining and immunofluorescent microscopy.

### Wounding Model

C57BL/6-Il17atm1Bcgen/J mice between 20-24 weeks of age were anesthetized with a mixture of 1.75L/m O2 and 2.5% of isoflurane and then received a 2-mm punch biopsy wound on one ear. Mice were individually housed and monitored daily. At 1-3 days post-wounding, the ears were harvested for staining and immunofluorescent microscopy.

### Epidermal Sheet Immunofluorescent Staining and Microscopy

Epidermal sheets were prepared as previously described (26). Briefly, ears were split in half and placed on ammonium thiocyanate solution (1x dPBS, 3.6% ammonium thiocyanate) dermis side down, and incubated for 15 minutes, after which the epidermis was peeled from the dermis. Epidermal sheets were floated on 1x dPBS for 5 minutes, then floated on 2 μg/ml anti-Vγ5 and anti-CCR6 (BioLegend, San Diego, CA) for 1 hour at 37°C with 5.0% CO_2_. Epidermal sheets were rinsed on 1x dPBS and mounted with Slowfade Gold Antifade Reagent with DAPI (Invitrogen, Waltham, MA). Slides were examined using an immunofluorescent microscope (Nikon DS-Qi2, Nikon, Tokyo, JPN). Expression of IL-17A (GFP), CCR6 (PE), and the γδTCR (APC) were examined at the site of wounding or psoriasis induction. Images were captured using NIS-Element D 4.1300 64 program (Melville, NY) at original magnification x200, and processed using Adobe Photoshop 2022 (San Jose, CA). Total epidermal γδ T cell number, as well as the number of epidermal γδ T cells that expressed CCR6, and/or IL-17 were quantified and then calculated as cells/mm^2^. In both the psoriasis and wounding experiments, a total of 7 regions were photographed per epidermal sheet per ear, accounting for both wounded and non-wounded ears, as well as Imiquimod-treated and untreated control ears. Each ear provided two epidermal sheets. In total 672 images were captured, analyzed and quantified between the Imiquimod and wounding models.

### sc-RNA Sequencing Data Processing and Analysis

Publicly available scRNA seq data was downloaded as fastq files from the NIH Gene Expression Omnibus (GEO) database (GSE149121) (20). Files were downloaded to the Linux terminal and transferred to the Cellranger Analysis Pipeline terminal. The study by Liu, et al. 2020 used single cell transcriptomics of CD45^+^ cells from mice treated with IMQ for 7 days (20). All fastq files were individually run through Cellranger Analysis Pipelines v6.1 (10x Genomics) to perform gene quantification and sequence alignment to the 10x Genomics mouse reference genome (mm10). Additionally, Cellranger was utilized to subsample experiment reads and produce an aggregated gene expression matrix. Once aggregation and structuring of the data was completed, Cellranger generated a cloupe file that was uploaded to the visualization software Loupe Browser v6.0 (10x Genomics) to be used for downstream analysis of scRNAseq data. scRNAseq data was then uploaded to Loupe Browser for further analysis. Epidermal γδ T cell populations were identified by their expression of *Tcrg-v5, Fcer1g* and *CD3.* Any contaminating non epidermal γδ T cell populations were eliminated based on their positive expression of *Cd4, Cd8, Tcrg-v4, Tcrg-v6, Krt5, Krt10, Cd207* or *Lyz1*. Once the epidermal γδ T cell clusters were identified, further analysis was performed by clustering the epidermal γδ T cells into subsets (CCR6^+^IMQ, CCR6^+^control, CCR6^−^IMQ, CCR6^−^control). Subsets were compared between IMQ treated and untreated control mice and differentially expressed genes (DEGs) were exported from Loupe Browser and submitted to Ingenuity Pathway Analysis (IPA) (Qiagen) for core analysis.

### Statistical Analysis

Unpaired Student t-tests and z-test were used to analyze T cell numbers and corresponding cytokine production. All statistical tests performed using Prism Graph Pad software (Dotmatics, Boston, MA). All findings are considered significant at *p* < 0.05. Log2-fold change in Loupe Browser was calculated by using the localized ratio of normalized mean gene UMI counts in each cluster relative to all other clusters.

## Results

### CCR6 is upregulated by a subset of epidermal γδ T cells upon anti-CD3 stimulation

Epidermal γδ T cells do not express CCR6 on the cell surface during homeostatic conditions (25). While resting epidermal γδ T cells do not express CCR6, we tested whether CCR6 is expressed upon activation. Epidermal γδ T cell lines were stimulated with or without anti-CD3 stimulation for 24 hours and CCR6 expression was examined by flow cytometry. Live, Vγ5 TCR^+^ cells were gated and analyzed for CCR6 expression. Upon stimulation with anti-CD3 for 24 hours, a subset of epidermal γδ T cells express CCR6 and CD25 (Fig. 1A). Of note is the role that may be played by IL-2 in the CCR6-expressing population.

**Figure 1.**
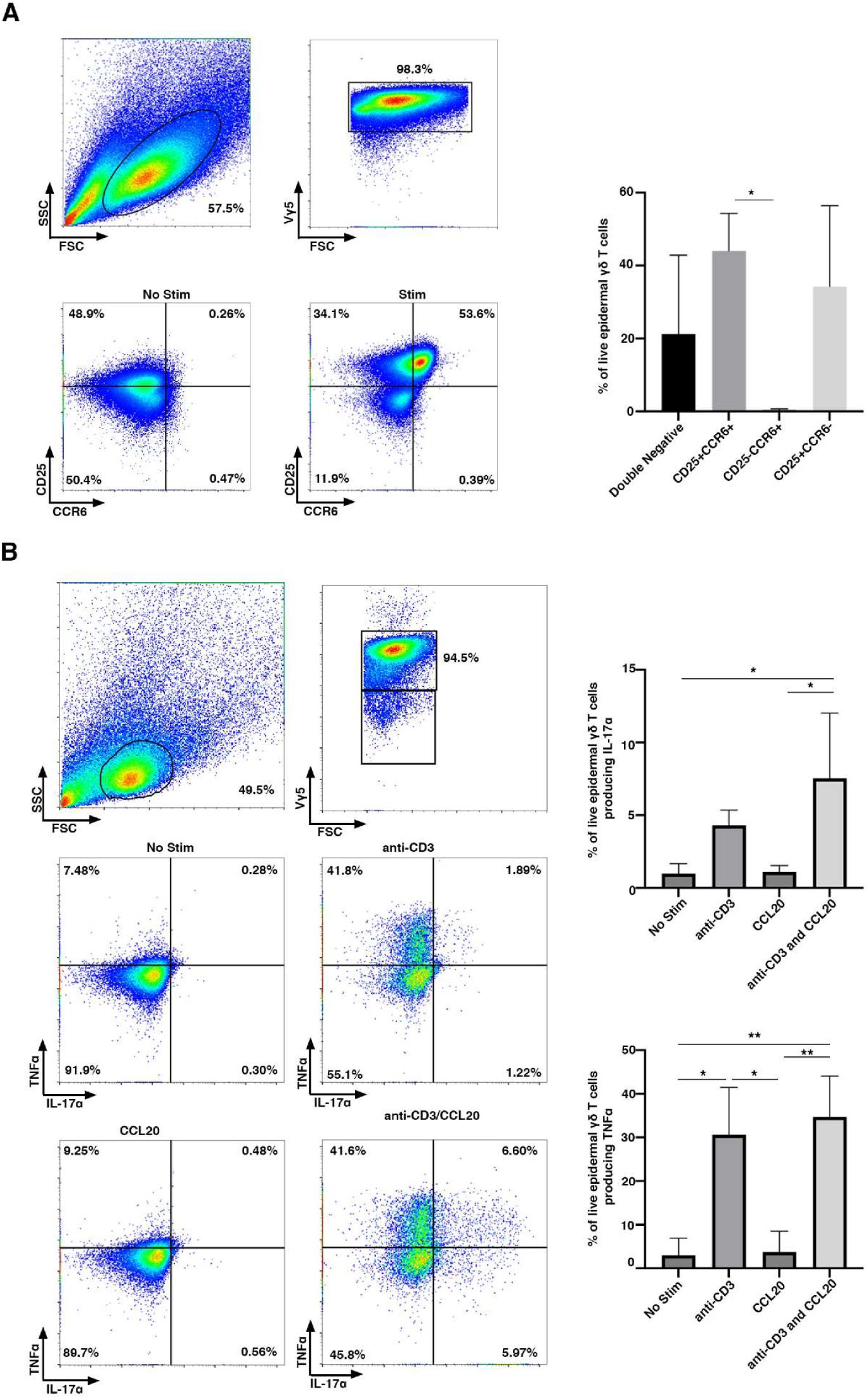
CCR6 is upregulated by a subset of CD25^+^ epidermal γδ T cells upon anti-CD3 stimulation. **(A)** Flow cytometric analysis of epidermal cells isolated from B6 mice, cultured for 9-15 weeks (> 95% epidermal γδ T cells), and stimulated with anti-CD3. Live, γδ TCR^+^ cells are gated and CCR6 and CD25 analyzed (n=3). **(B)** Flow cytometric analysis of epidermal Vγ5 T cells stimulated in the presence or absence of anti-CD3 and/or CCL20 for 6 hours. Live, Vγ5^+^ T cells are gated and TNF-α and IL-17A analyzed (n=3). Data represents the mean +/− SD. \**p* < .05 and ** *p* < .01

### CCL20 increases IL-17A, but not TNF-α production by activated epidermal γδ T cells

Previous studies show that CCR6^+^ dermal Vγ4 T cells secrete IL-17A in response to skin damage and during psoriasis (13, 16, 27). Lower levels of IL-17A have also been detected by epidermal γδ T cells during wound repair and contact hypersensitivity, but it is unknown if this is regulated by CCL20 and whether this is a distinct population (15, 16). To determine whether IL-17A and TNF-α production by epidermal γδ T cells is augmented by the CCR6 ligand, CCL20, we examined epidermal γδ T cells post-stimulation with CCL20 and/or anti-CD3. There is not a significant increase in epidermal γδ T cells producing IL-17A post-stimulation with anti-CD3, but there is a significant increase in epidermal γδ T cells producing TNF-α (Fig. 1B). CCL20 alone does not increase the percent of epidermal γδ T cells producing either IL-17A or TNF-α. Furthermore, CCL20 administered with anti-CD3 stimulation does not induce more epidermal γδ T cells to produce TNF-α than anti-CD3 alone (Fig. 1B). However, CCL20 administered with anti-CD3 stimulation significantly increases the percent of epidermal γδ T cells producing IL-17A. Interestingly, upon the addition of anti-CD3 and CCL20, there are four functional subsets of epidermal γδ T cells: TNF-α^−^/ IL-17A^−^, TNF-α^+^/IL-17A^−^, TNF-α^−^/IL-17A^+^ and TNF-α^+^/ IL-17A^+^. This challenges previous studies suggesting that epidermal γδ T cells are preprogrammed away from a Tγδ17 fate.

### Epidermal γδ T cells expressing CCR6 exhibit a unique gene expression profile

To examine gene expression by CCR6^+^ epidermal γδ T cells during psoriasis, publicly available scRNAseq data from Liu et. al was reanalyzed with a focus on CCR6^+^ or CCR6^−^ epidermal γδ T cells (20). In this study RNA was isolated from C57BL/6J mouse skin treated with or without 5% Imiquimod for seven days and scRNAseq was performed. The authors report a cluster of epidermal γδ T cells in a t-distributed stochastic neighbor embedding (t-SNE) plot indicating a broad population with a potential for subsets of epidermal γδ T cells with differing gene expression. To specifically analyze epidermal γδ T cells, target cells were sorted from the overall cell count using Loupe Browser based on well-established marker genes (*Cd3^+,^ Tcrg-v5^+^, Trdc^+^, Cd4^−^, Cd8^−^*). Minor populations of contaminant keratinocytes and Langerhans cells were excluded by removing cells exhibiting high levels of *Krt5*, *Krt10* and *Krt14* for the keratinocytes and high levels of *Cd207* for the Langerhans cells, yielding a total of 236 epidermal γδ T cells for analysis (Fig. 2A). These exclusions produced three distinct epidermal γδ T cell populations (labeled cluster 1-3 in Fig. 2A).

**Figure 2.**
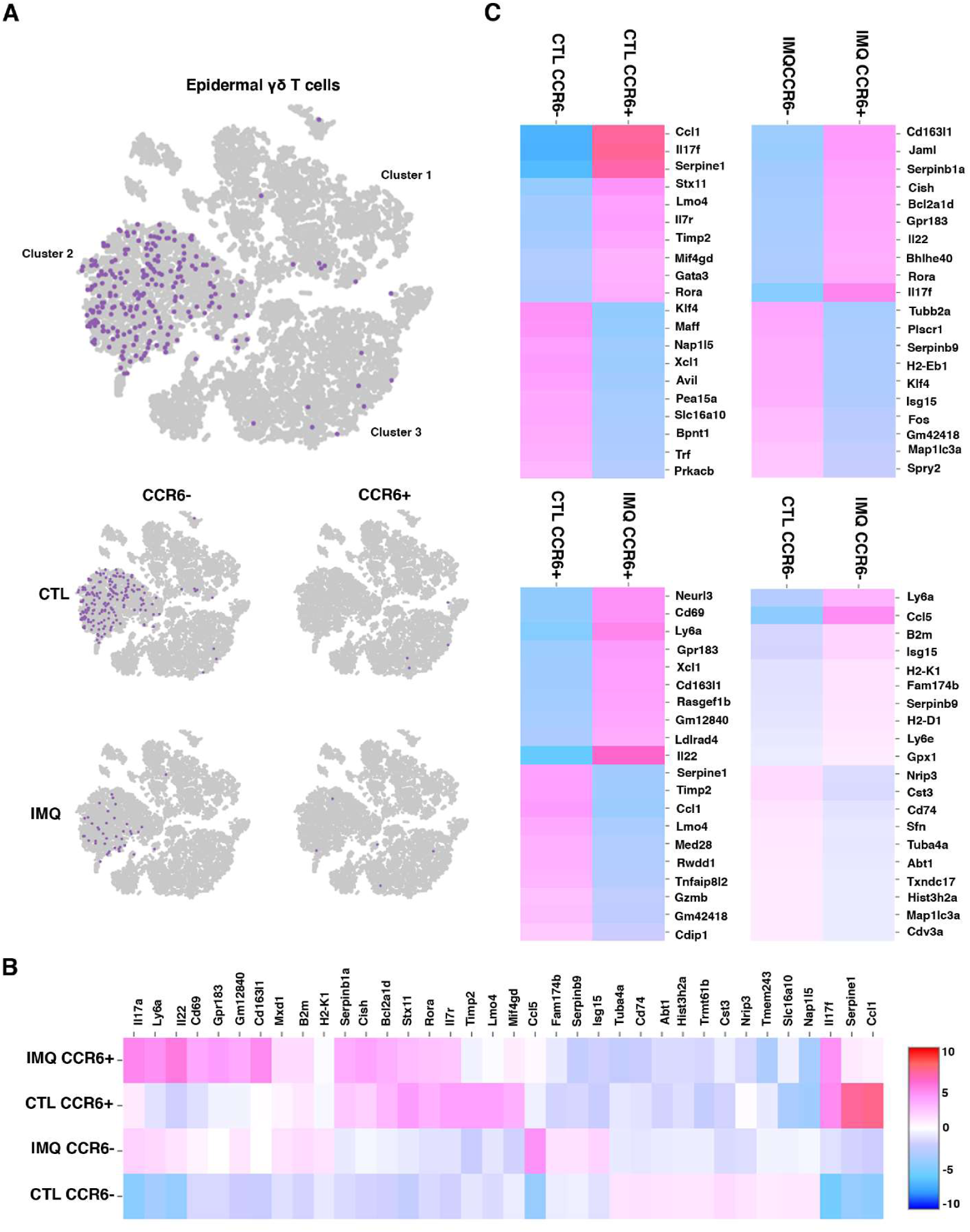
CCR6^+^ epidermal γδ T cells exhibit a Tγδ17 gene expression profile. Using publicly available scRNA-Seq data (20) the total skin cell population (n=18,040) was filtered to epidermal γδ T cells (n=236). **(A)** UMAP plot showing epidermal γδ T cells can be identified in three distinct clusters. Based on treatment group and CCR6 expression epidermal γδ T cells were clustered further into CCR6^+^IMQ, CCR6^+^control, CCR6^−^IMQ, CCR6^−^control. In the treatment group, the ratio of CCR6^+^ to CCR6^−^ cells increased from 1/48 in the control group to 1/9. **(B)** Differential gene expression between 4 different epidermal γδ T cell populations within treatment groups displayed in a heatmap **(C)** Dual heatmap rendering of differential gene expression between specific epidermal γδ T cell groups.

Epidermal γδ T cells were further clustered based on treatment group and CCR6 expression (CCR6^+^IMQ, CCR6^+^control, CCR6^−^IMQ, CCR6^−^control) (Fig. 2A). Prior to IMQ treatment, CCR6^−^ epidermal γδ T cells are represented in clusters 1, 2 and 3. However, post-IMQ treatment the CCR6^−^ epidermal γδ T cells have predominantly centralized to cluster 2. Pre-IMQ treatment CCR6^+^ epidermal γδ T cells are represented in cluster 3, while post-treatment the CCR6^+^ epidermal γδ T cells are represented in both cluster 2 and cluster 3. A locally distinguishing differential gene expression (DGE) analysis was run between the CCR6+ and CCR6-epidermal γδ T cell populations with and without IMQ treatment and a heat map was generated showcasing the top 10 significantly upregulated genes per cluster based on the logarithmic fold change of gene expression between each group during paired comparison (Fig. 2B).

Interestingly, both CCR6^+^ subsets differentially express the transcription factor *Rora*, which regulates both CCR6 and IL17 expression (Fig. 2B) (28). To better characterize each individual epidermal γδ T cell population we analyzed DEGs between paired subsets. Comparing the CCR6^+^control and CCR6^−^control groups, we observe that the CCR6^+^control gene profile is more inflammatory, favoring immune cell infiltration (*Ccl1, Serpine1, Il17f*), whereas the CCR6^−^ control group does not exhibit the upregulation of any DEGs above 5-fold. However, when examining gene expression changes exceeding 4-fold, the CCR6^−^control subset demonstrates increased expression of genes related to mitotic cell cycle, DNA binding, and cell motility (*Maff, Klf4*) (Fig. 2C top left). Next, in the comparison between CCR6^+^IMQ and CCR6^−^IMQ treatment groups, the CCR6^+^IMQ subset exhibits a Tγδ17 inflammatory profile, with a >5-fold upregulation of *Il17f*, along with lower level increases in *Rora*, *IL22*, and *Jaml*. While the CCR6^−^IMQ subset experiences a >3-fold upregulation in genes associated with cytokinesis, proliferation and cell motility (*Tubb2a, Fos, Plscr1*) (Fig. 2C top right).

In the comparison among CCR6^+^ groups, the CCR6^+^IMQ subset consistently exhibits a pro-inflammatory profile compared to all other subsets, with a greater than 5-fold change in *Il22* expression (Fig. 2C bottom left). On the other hand, the CCR6^+^control subset still shows a >3-fold upregulation in genes associated with immune cell infiltration (*Ccl1, Serpine1*). Lastly, the comparison among CCR6^−^ subsets indicate minimal gene upregulation except for potential immune cell recruitment (*Ccl5*) in the CCR6^−^IMQ subset (Fig. 2C bottom right). Our results suggest that CCR6 expressing epidermal γδ T cells in psoriasis exhibit a skewed IL17 focused response.

### Psoriasis increases MYC pathway signaling by CCR6^+^ epidermal γδ T cells

IPA core analysis was performed to further elucidate the biological processes and molecular mechanisms that differentiate CCR6^+^ epidermal γδ T cell subsets in psoriasis. Top differentially expressed genes were clustered into canonical pathways using IPA Knowledge Base platform (Fig. 3A). Comparison of CCR6^+^IMQ to CCR6^−^IMQ subsets revealed Eukaryotic Translation Elongation, Eukaryotic Translation Termination, Response of EIF2AK4 (GCN2) to Amino Acid Deficiency, SRP-dependent Cotranslational Protein Targeting to Membrane, Eukaryotic Translation Initiation, Nonsense-Mediated Decay (NMD), Selenoamino Acid Metabolism, EIF2 Signaling and Major Pathway of rRNA Processing in the Nucleolus and Cytosol as the significantly upregulated pathways of CCR6^+^IMQ subsets in psoriasis when compared to CCR6^−^IMQ subsets in psoriasis (Fig. 3A). Examination of significantly differentiated upstream regulators between these subsets revealed that the MYC pathway is significantly upregulated within the population of CCR6^+^IMQ epidermal γδ T cells when compared to CCR6^−^IMQ epidermal γδ T cells (Fig. 3B). The MYC pathway is predicted by the IPA Knowledge Base to be linked with activation of the transcription factor *Rora* (Fig. 3B). Analysis of the *Rora* downstream pathway shows that upregulation of *Rora* by CCR6^+^IMQ epidermal γδ T cells directly activates the *Il17a* and *Il17f* expression found within CCR6^+^IMQ epidermal γδ T cell subsets. Furthermore, this downstream pathway activation by *Rora* is predicted by IPA Knowledge Base to lead to activation of *Il22* as well as the CCL20/CCR6 axis in CCR6^+^IMQ epidermal γδ T cell subsets (Fig. 3C). Our results suggest that the activation of the MYC pathway within CCR6^+^IMQ epidermal γδ T cell subsets contributes to the increased Tγδ17 cytokine profile observed within CCR6^+^IMQ epidermal γδ T cell subsets.

**Figure 3.**
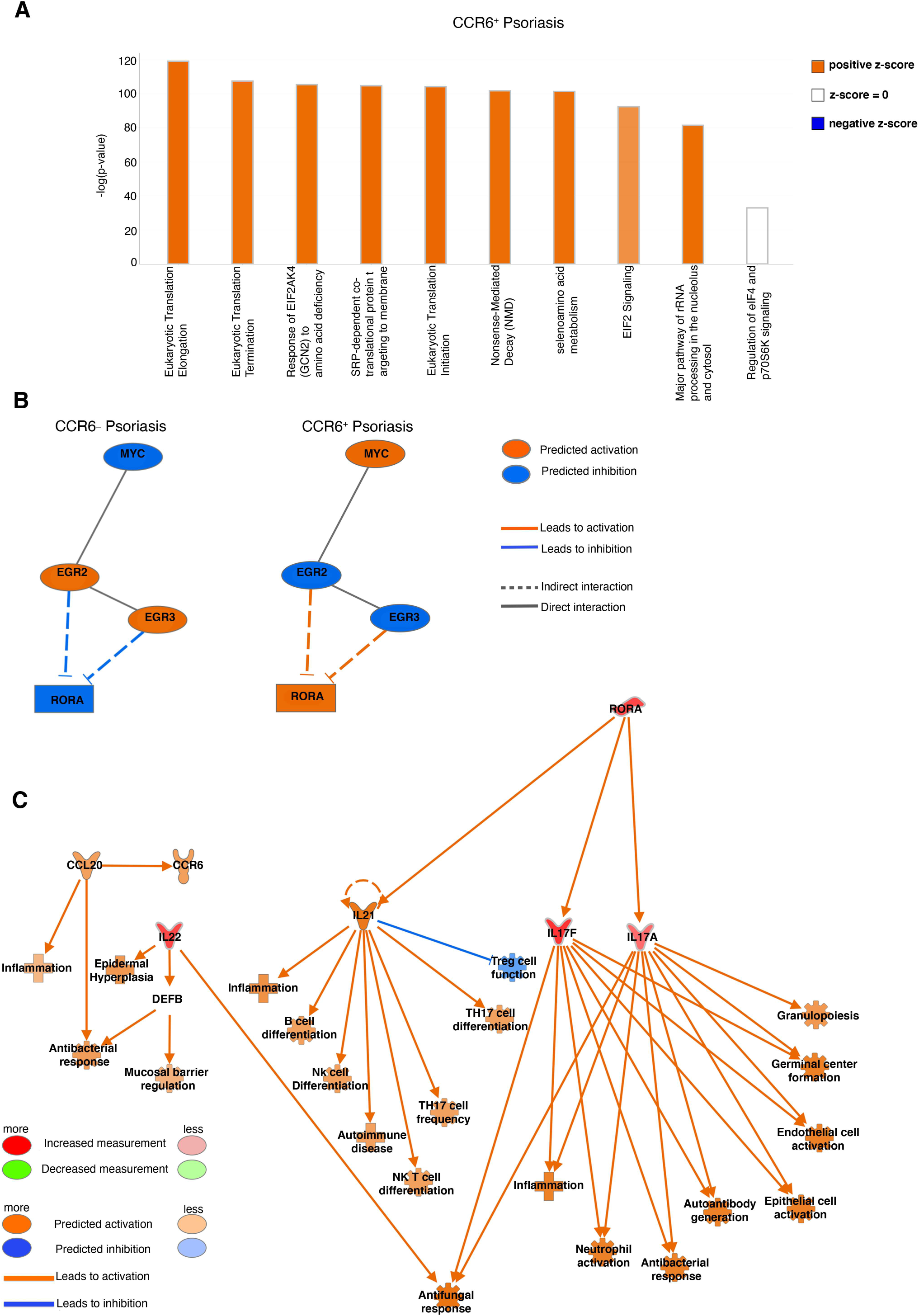
Canonical pathway analysis of differentially expressed genes from CCR6^+^ and CCR6^−^ epidermal γδ T cells during IMQ-induced psoriasis identify Myc pathway. **(A)** Top differentially expressed genes between CCR6^+^IMQ and CCR6^−^IMQ from publicly available scRNA sequencing data (20) were clustered into canonical pathways using IPA Knowledge Base platform. Positive z-score (orange) shows pathway activation. A negative z-score (blue) shows the pathway is inhibited. No z-score (white) shows the pathway is neither activated nor inhibited. Bars are arranged by statistical significance. **(B)** Network analysis identifies upstream regulator Myc (orange indicating activation), with IPA Knowledge Base predicting a linked activation with Rora. **(C)** Network downstream analysis of Rora pathway with orange predicting activation, blue predicting inhibition, and red predicting an increased measurement between subsets (CCR6^+^IMQ vs. CCR6^−^IMQ epidermal γδ T cells).

### Psoriasis increases IL-17 and CCR6 production by epidermal γδ T cells

To validate the scRNAseq findings that CCR6^+^ epidermal γδ T cells produce IL-17A during psoriasis, we examined IL-17A production *in vivo* using IL-17A GFP reporter mice. To establish when CCR6 upregulation occurs post-IMQ treatment, a pilot experiment was performed using C57BL/6J wild type mice. Pilot results indicate that CCR6 expression by epidermal γδ T cells peaks after two days of IMQ treatment. Thus, in our studies, mice received IMQ treatment for two days prior to analysis. This timepoint is similar to previous studies in wound healing models when epidermal γδ T cell activation was observed within 6 hours of wounding and function persists for at least two days (17, 29). Epidermal sheets were costained for Vγ5, and CCR6, while IL-17A was detected with GFP, and epidermal γδ T cells quantified per mm^2^ (Fig. 4A). During IMQ-induced psoriasis, there are more IL-17A-producing epidermal γδ T cells than controls (Fig. 4A, B), but this increase is just short of reaching significance (p=0.075) (Fig. 4B). Similarly, there are more CCR6-expressing epidermal γδ T cells upon IMQ treatment (Fig. 4A, 4B). While CCR6^+^IL-17A^+^ epidermal γδ T cells were easily identified in IMQ treated mice, the increases that did not reach significance. Similarly, no significant differences were found in the percentage of CCR6^+^ epidermal γδ T cells that were also IL-17A^+^ (Fig. 4A, 4B). However, given that the epidermal γδ T cells express both CCR6 and IL17A in a time-dependent manner, CCR6 may be upregulated prior to IL-17A.

**Figure 4.**
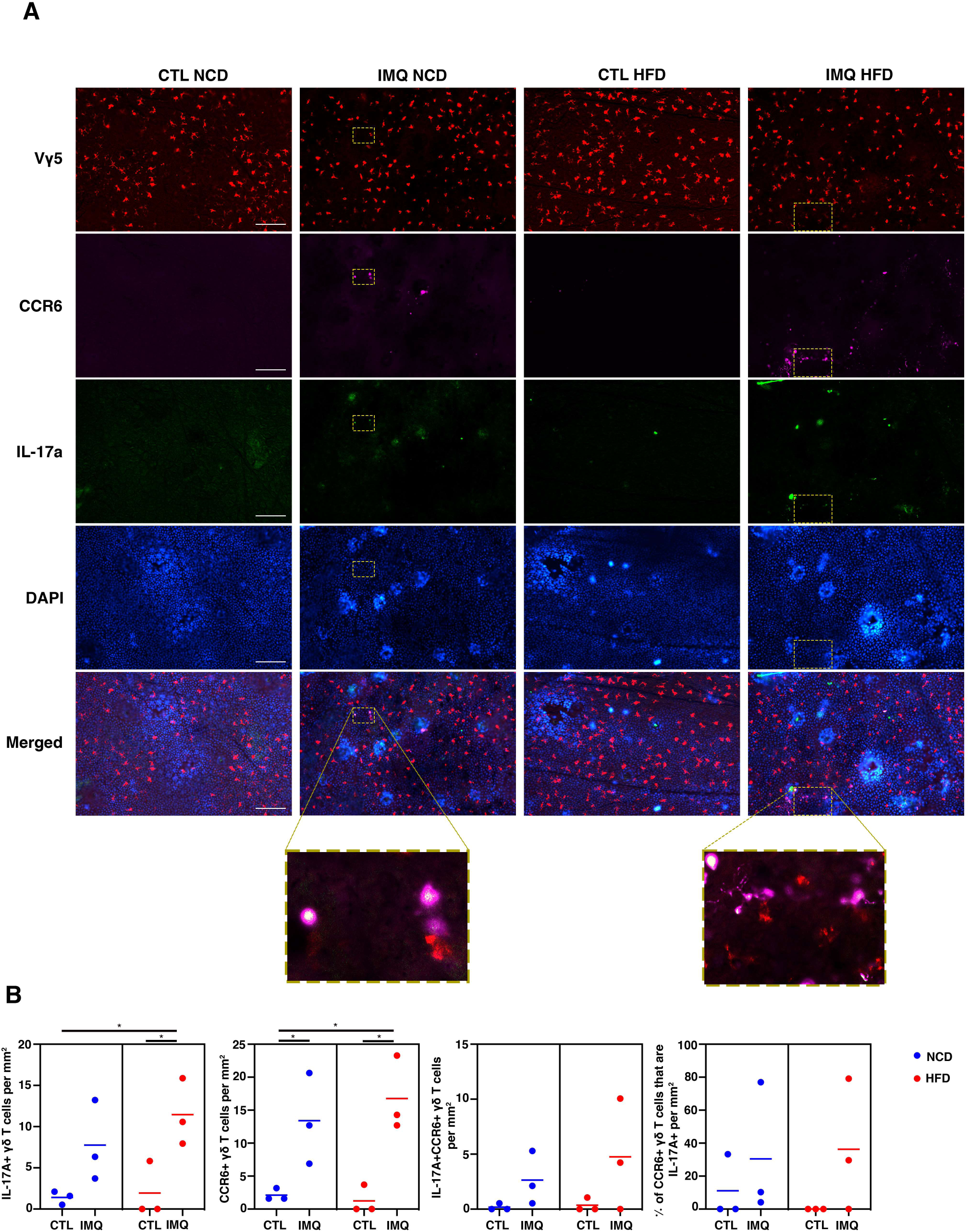
IMQ-induced psoriasis increases IL-17A and CCR6 expression by epidermal γδ T cells, while obesity does not further increase expression. **(A)** Representative immunofluorescent images of epidermal sheets from C57BL/6-Il17atm1Bcgen/J mice treated with and without IMQ for two days. Scale bars, 100 μm. **(B)** Quantification of IL-17A^+^, CCR6^+^, and IL-17A^+^CCR6^+^ epidermal γδ T cells with and without IMQ treatment (n=3 mice/group). A minimum of seven fields of view for each mouse were used for analysis and then averaged to form one data point. * *p* < 0.05

### IL-17 and CCR6 upregulation by epidermal γδ T cells in obese and lean mice is similar in psoriasis

To determine whether there is an impact of obesity on CCR6 expression or IL-17 production by epidermal γδ T cells in psoriasis-like inflammation, IL-17A GFP reporter mice were fed either a NCD or HFD for 12-16 weeks and then received IMQ treatment for two days. There is a significant increase in IL-17 production by epidermal γδ T cells within the IMQ group when compared to the control group in mice fed a HFD (Fig. 4A, 4B). This is now significant where it was only reaching significance in the lean control group suggesting a subtle effect of obesity on IL-17 production by epidermal γδ T cells. Upregulation of CCR6 by epidermal γδ T cells in psoriasis occurs in both NCD and HFD-fed mice to a similar degree, suggesting obesity does not exacerbate CCR6 expression at this timepoint of IMQ-induced psoriasis onset (Fig. 4A, B).

### CCR6 is upregulated within 1 day and downregulated by 3 days post-wounding

To determine how CCR6 is regulated by epidermal γδ T cells in response to wounding, we performed a time course. Epidermal γδ T cells are known to become activated and produce growth factors and cytokines for the first two days post wounding. Thus, we examined CCR6 expression on epidermal γδ T cells at days 0, 1, 2 and 3 post wounding. As expected, there is little to no CCR6 expression by epidermal γδ T cells in non-wounded control skin (Fig. 5A,B). However, the number of CCR6 expressing cells increases significantly 1 day post wounding, with CCR6 expression returning to nonwounded control levels 3 days post wounding (Fig. 5A,B). Overall, this data indicates that CCR6 expression by epidermal γδ T cells is early and temporal during wound repair instead of being constitutive as in dermal γδ T cells.

**Figure 5.**
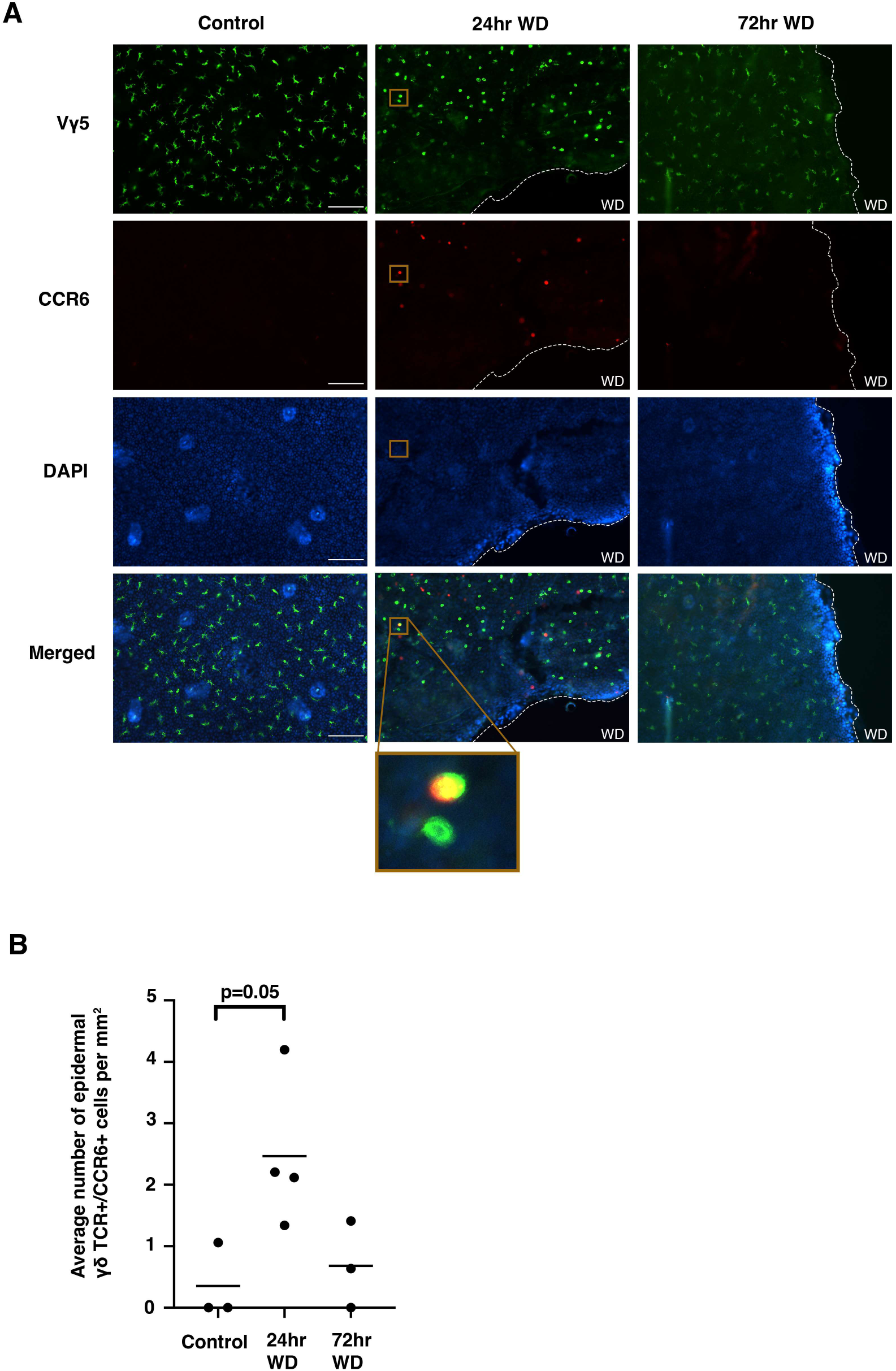
CCR6 is upregulated within the first 24 hrs. post-wounding and downregulated by 72 hrs. post-wounding. **(A)** Representative immunofluorescent images of epidermal sheets from B6 mice at various timepoints post wounding. Wound site indicated with dotted line. Scale bars,100 μm. **(B)** Quantification of CCR6^+^ epidermal γδ T cells at different timepoints post wounding (n= 3-4 mice/group). A minimum of seven fields of view for each mouse were used for analysis and then averaged to form one data point. *p* = .05

### IL-17A-expressing epidermal γδ T cells are significantly increased at the wound site

CCR6 and IL-17 production by epidermal γδ T cells were examined in wounded and non-wounded IL-17A reporter mice. IL-17A-producing epidermal γδ T cells were increased in wounded mice as compared to their non-wounded counterparts. CCR6-expressing epidermal γδ T cells from wounded NCD mice were increased in two of the three wounded mice as compared to nonwounded mice (Fig. 6A, 6B). 20% of the CCR6^+^ epidermal γδ T cells express IL-17A in wounded mice compared to 0% in non-wounded mice. Together this data suggests that wounding induces epidermal γδ T cells to upregulate IL-17 and this occurs on a proportion of CCR6^+^ cells.

**Figure 6.**
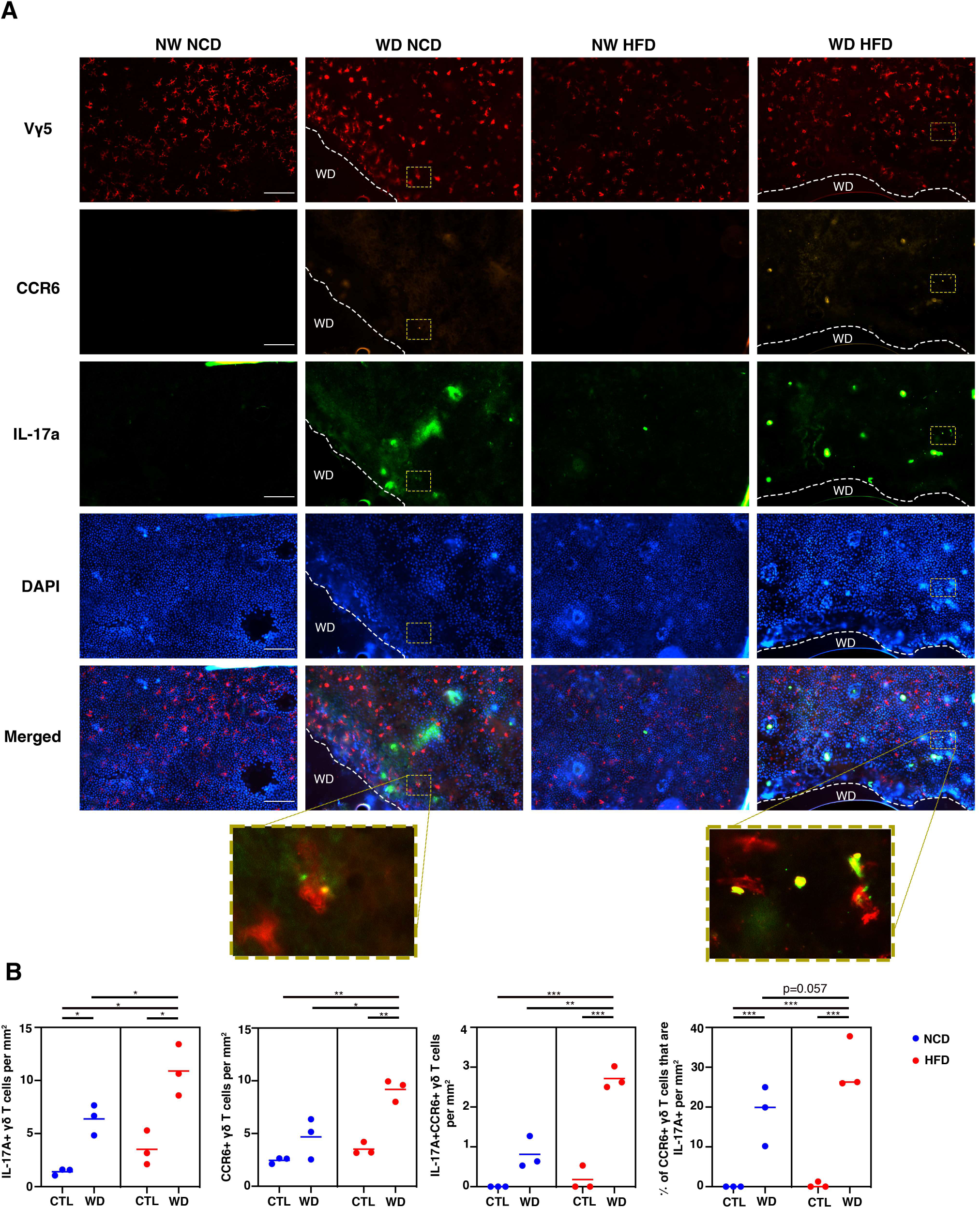
IL-17A and CCR6 expression by epidermal γδ T cells is significantly elevated in wounded mice, while obesity increases IL-17A production and CCR6 expression by epidermal γδ T cells at the wound site. **(A)** Representative immunofluorescent images of epidermal sheets from C57BL/6-Il17atm1Bcgen/J mice one day post wounding. Wound site indicated with dotted line. Scale bars, 100 μm. **(B)** Quantification of IL-17A^+^, CCR6^+^, and IL-17A^+^CCR6^+^ epidermal γδ T cells with and without wounding (n=3 mice/group). A minimum of seven fields of view for each mouse were used for analysis and then averaged to form one data point. * *p* < 0.05, ** *p* < .01, *** *p* < .001

### Obesity increases IL-17A production and CCR6 expression by epidermal γδ T cells at the wound site

Previous studies have shown that obesity alters epidermal γδ T cell number and function during wound repair (8, 14). To determine if there is a shift in epidermal γδ T cell function toward a Tγδ17 phenotype in obesity IL-17A GFP reporter mice were fed either a NCD or HFD for 12-16 weeks and then were wounded for 1 day prior to analysis. Obese mice exhibit an increase in IL-17A-producing epidermal γδ T cells in wounded vs non-wounded mice (Fig. 6B). In addition, at the wound site, obese mice exhibit significantly elevated numbers of IL-17A-producing epidermal γδ T cells as compared to their NCD counterparts (Fig. 6B). CCR6 expression by epidermal γδ T cells at the wound site is also significantly increased in obese mice as compared to lean mice (Fig. 6A, 6B). The number of CCR6^+^ epidermal γδ T cells that simultaneously express IL-17A is also higher in obese mice upon wounding and as compared to lean wounded mice (Fig. 6B). There is also an increase in the percentage of CCR6^+^ epidermal γδ T cells concurrently expressing IL-17A, which is nearly significant in the wounded obese group as compared to the wounded lean group (p=0.057) (Fig. 6B). Together these data show that obesity increases Tγδ17 epidermal γδ T cells.

## Discussion

Epidermal γδ T cells exhibit a variety of functional responses including Tγδ1, Tγδ2, and Tγδ17 for roles in wound repair, tumor cytolysis and contact hypersensitivity (15, 17, 30). These functions are regulated via costimulation through receptors such as JAML and CD100 in wound repair and cytokine reception such as IL-1β in contact hypersensitivity (16, 31). Here we suggest that there are functional subsets of epidermal γδ T cells with skewed abilities and that these subsets can be increased by environmental factors such as obesity. Previously, epidermal γδ T cells with IFN-γ and IL-17 secreting abilities were identified, but there are currently no markers to further define these cells and to define whether these are specific subsets or cellular plasticity (16). We show that the expression of CCR6 upon activation defines a subset of epidermal γδ T cells with Tγδ17 functional capabilities. Thus, chemokines such as CCL20 can direct the function of a distinct epidermal γδ T cell subset during activation.

It is possible that identifying subsets of epidermal γδ T cells has been difficult because the markers are more easily observed during activation. Particularly for the CCR6^+^ subset, T cell activation is required for the upregulation of CCR6 and then chemokine reception is required for IL-17A production. Our data suggest that there is a temporal requirement for TCR signaling, followed by CCR6 upregulation and CCL20 reception to get IL-17A production. We reveal an association between the Tγδ17 profile observed in CCR6^+^ epidermal γδ T cells during psoriasis and the upstream transcription factor Myc. Myc is an early response gene in T cell activation (32). Expression of Myc is regulated by TCR signal strength and cytokine reception such as IL-2 (33). This correlates well with our finding that CCR6^+^ epidermal γδ T cells also express CD25. Further, we find that Myc activation is positively associated with RORα, primarily attributed to the inhibition of early growth response protein 2 (EGR2). EGR2 deficiency in CD4 T cells leads to an increase in IL-17 expression (34). Furthermore, it has been established that the overexpression of Myc by γδ NKT cells results in EGR2 deficiency (35). Our results are consistent with published studies which indicate RORα directly regulates the expression of both CCR6 and IL-17 (36). When considering the known interaction between RORα, EGR2, and Myc, our results suggest that the pathway involving Myc, EGR2, and RORα may serve as a promising focus for understanding the underlying mechanisms behind the expression of the Tγδ17 profile in CCR6^+^ epidermal γδ T cells during skin inflammation.

Obesity increases the number of CCR6 and IL-17-expressing epidermal γδ T cells during the early stages of wound repair, but not during IMQ-induced psoriasis. It is possible that there is a set number of epidermal γδ T cells with preprogrammed Tγδ17 function and that IMQ-induced psoriasis activates the entire subset. Thus, obesity would not induce an additional increase, while wounding only induces some of the Tγδ17 epidermal T cells and obesity further increases the number of activated Tγδ17 epidermal T cells. Previous research has demonstrated that obesity not only upregulates IL-17 but also boosts the production of CCL20 (24, 37). This dysregulation of the epidermis corresponds to our previous findings that show the cellular composition and organization of the epidermis is altered in HFD and db/db mouse models of obesity due to hyperglycemia and chronic inflammation. In diabetes and obesity, T cell and keratinocyte numbers and tissue repair functions are compromised. Epidermal γδ T cell dysfunction plays a role in the reduced number and increased differentiation of keratinocytes in both models (8, 14). Another theory is that obesity does not increase IL-17 and CCR6-expressing epidermal γδ T cells during psoriasis because of the experimental timeline we used. During IMQ-induced psoriasis, IL-17 production is evident in skin resident T cell populations, and continues to rise throughout the 7 day IMQ treatment (20). For this study we chose the timepoint in which we observed the most CCR6^+^ epidermal γδ T cells in lean mice, but obesity may alter that timepoint.

CCR6 has been associated with numerous diseases including psoriasis and is a target for therapeutic intervention, but CCR6 targeting drugs have not yet been approved by the FDA (38, 39). Cells including dermal γδ T cells utilize CCR6 to traffic to the epidermis during inflammation (21, 40). Thus, drugs that block the function of CCR6 or interaction with CCL20 would reduce the recruitment of IL-17-producing T cells. Epidermal γδ T cells normally reside in the basal layer between keratinocytes and are not known for migrating within or outside of the epidermis (17, 41). Here we have identified a clear subset of epidermal γδ T cells that upregulate CCR6 and thus would also be targeted with CCR6-specific therapeutics. It is now clear that CCR6^+^ epidermal γδ T cells contribute to Tγδ17-associated responses in psoriasis and wound healing, challenging previous assumptions that other dermal and infiltrating cell types were the only IL-17-producers (13, 25). Further, this data showcases the impact of obesity on epidermal γδ T cell subsets and function in inflammatory settings.

## Abbreviations

IMQ: Imiquimod
ILC: innate lymphoid cell
MAIT: mucosal-associated invariant T
TCR: T cell receptor
scRNAseq: single-cell RNA sequencing
HFD: high fat diet
NCD: normal chow diet
DEG: differentially expressed gene
DC: dendritic cell
NK: natural killer cell
NMD: nonsense-mediated decay
WD: wounded
NKT: natural killer T cell.

## Acknowledgements

The authors thank the Jameson lab members for manuscript review.

